# Functional Interaction of SKA and NDC80 Complexes at Kinetochores Promoting Anaphase Onset in Mitosis

**DOI:** 10.64898/2026.05.22.727258

**Authors:** John R Daum, Natalia Romek, Gary J Gorbsky

## Abstract

The kinetochore and spindle complex (SKA) and NDC80 complexes are essential kinetochore elements that ensure highly accurate chromosome segregation and successful progression through mitosis. The SKA heterodimer complex consists of SKA1, SKA2, and SKA3 subunits, and the NDC80 complex contains NDC80, NUF2, SPC24, and SPC25 subunits. Through live cell fluorescence timelapse imaging assays and expression of RNAi-resistant SKA3 constructs, we rescue SKA complex function in cells lacking endogenous SKA3. These assays reveal a critical span within SKA3’s C-terminus required for successful mitotic progression. Structural protein modeling shows that this span encompasses the majority of a roughly 40 amino acid SKA3 C-terminal structural element that promotes interaction with the coiled-coil NDC80 and NUF2 subunits of the NDC80 complex. Thus, although spindle and kinetochore concentration of the SKA complex is mediated in part by the tubulin and tip-tracking capabilities provided by the SKA1 component of the SKA complex, transition from metaphase to anaphase requires the contribution of SKA3’s C-terminal structural interface to mediate interaction between the SKA and NDC80 complexes.

**Significance Statement:** Accurate chromosome segregation is essential for genomic stability, and its failure underlies developmental defects, and cancer. The kinetochore-microtubule interface, where the SKA and NDC80 complexes converge, is central, yet the of these two complexes is incompletely defined. We identify a critical segment within the C-terminus of SKA3 required for the metaphase-to-anaphase transition. Using live-cell imaging with RNAi-resistant rescue constructs and structural modeling, we demonstrate that this region mediates SKA engagement with the coiled-coil domains of NDC80 and NUF2. Our findings establish that a small segment of SKA3’s C-terminus provides an essential physical bridge between the SKA and NDC80 complexes. This work refines understanding kinetochore-microtubule interaction, functionally identifying a discrete contact whose disruption may be relevant to chromosomal instability in disease.

## Introduction

In a single day an estimated 330 billion cell divisions occur within the adult human body, an average rate roughly equivalent to 4 million per second (1). Progression through mitosis, a series of defined phases which enable these divisions, requires a resilient system to facilitate the interaction of paired replicated chromosomes with the microtubule-based mitotic spindle. This ensures the accurate distribution of duplicated chromosomes to newly forming daughter cells. A component of this system is SKA, the <u>S</u>pindle and <u>K</u>inetochore <u>A</u>ssociated complex (2–6). In metazoans SKA associates with nascent mitotic spindle microtubules at prophase and, post nuclear envelope breakdown, with kinetochores during prometaphase. Maximal kinetochore-localized concentrations are reached at metaphase, decrease during anaphase, and are lost at telophase when SKA is excluded from nuclei (5). The SKA complex is a W-shaped heterotrimeric assembly, consisting of 2 copies each of the SKA1, SKA2, and SKA3 subunits. SKA1 and SKA3 N-terminal domains and essentially the entire SKA2 subunit intertwine to assemble the complex (7). Enabled by interaction with binding partners and post translational modifications, SKA1 and SKA3 subunits support SKA’s regulatory functions.

Human SKA1 possesses a C-terminal tail that extends from the intertwined junction of the SKA subunits. Comprised of roughly the terminal 123 amino acids, SKA1’s extreme C-terminus is largely a globular domain encompassing residues 133-255 of the protein sequence. The N-terminal 1-91 amino acids have been shown to support SKA complex formation while the central 92-132 amino acids likely comprise a flexible bridge between the intertwined subunits and SKA1’s globular domain. The globular domain binds microtubules and the SKA complex’s concentration at kinetochores is partially dependent upon it, however SKA will accumulate at kinetochores in the absence of the SKA1 c-terminal globular domain or in the absence of microtubules (5, 8, 9). SKA1’s C-terminal domain is also responsible for kinetochore recruitment of protein phosphatase 1 (PP1), promoting timely anaphase onset and successful mitotic progression (8). HeLa cells entering mitosis without endogenous SKA1 but expressing a mutant SKA1 lacking its C-terminal globular domain delay or arrest at metaphase and frequently undergo cohesion fatigue, a process of asynchronous sister chromatid separation induced by sustained pulling forces of the mitotic spindle (8, 9). This phenotype is consistent amongst HeLa cells lacking functional SKA complex, as demonstrated by RNA interference (RNAi) targeting individual or multiple SKA complex subunits (2–6).

In addition to binding microtubules and recruiting protein phosphatase 1 to kinetochores, the SKA complex interacts with the NDC80 complex, an association that’s required for normal mitotic progression. The heterotetrametric NDC80 complex is composed of one copy each of NDC80 (also known as HEC1 in mammalian nomenclature), NUF2, SPC24, and SPC25 proteins. A dumbbell shaped molecular machine, the complex links microtubules of the mitotic spindle to the centromere-proximal inner kinetochore from prometaphase through anaphase. Its primary functions are to track with microtubule plus ends, enabling prometaphase chromosome congression, and to form and coordinate load bearing, stable microtubule plus-end bipolar attachments. Achieved at metaphase, these stable bipolar attachments between the mitotic spindle and sister chromatids ensure duplicated chromosomes will be distributed to opposite spindle poles during anaphase. NDC80 complex subunits NDC80 and NUF2 possess N-terminal calponin homology (CH) domains that form spindle-facing globular heads and interact with tubulin subunits of microtubules. The coiled C-termini of NDC80 and NUF2 bind the N-termini of SPC24 and SPC25 subunits. A tightly wound heterodimer of the centromere-facing end of the complex consists of the C-termini of SPC24 and SPC25, ending in a singular globular head that anchors the NDC80 complex to kinetochores (10).

Prior to achieving stable end-on kinetochore attachments with microtubules at metaphase, the NDC80 complex is essential for spindle checkpoint signaling. The spindle checkpoint is a kinetochore-driven error correction system promoting the dissolution of inappropriate kinetochore-microtubule interactions. This prevents anaphase onset prior to the generation of appropriate bipolar interactions known as amphitelic attachments (11, 12). Without stable and fully matured amphitelic interactions, opposing poleward forces creating tension across paired kinetochores of replicated chromosomes are not realized or maintained. Under these conditions the checkpoint kinase MPS1 promotes continued kinetochore synthesis of the mitotic checkpoint complex (MCC) and along with another checkpoint kinase, Aurora B kinase, works to destabilize inappropriate microtubule interactions so that amphitelic attachments will dominate. While the active spindle checkpoint delays mitotic progression and promotes error correction, increasing amphitelic microtubule-NDC80 complex interactions strengthen poleward forces and alter kinetochore structure. As the kinetochore and NDC80 complex change shape due to increasing spindle forces, mitotic checkpoint kinase phosphorylation of substrates is reduced by physical separation of kinase and substrate and by loss of kinetochore concentration of these kinases and substrates. Dephosphorylation of mitotic checkpoint kinases and their substrates reduces the production of the MCC and promotes spindle checkpoint silencing. SKA’s recruitment of PP1 through SKA1’s globular C-terminal domain likely contributes to this targeted dephosphorylation of substrates.

Akin to SKA1, SKA3 possesses a C-terminal tail that extends from its junction with the SKA1/2 subunits. Analysis of its amino acid sequence suggests that it is largely unstructured and disordered. Utilizing cross-linking mass spectrophotometry experiments, Helgeson et al. found that the SKA complex likely binds the NDC80 complex through direct interactions via this unstructured C-terminal region of SKA3, finding crosslinks between SKA3’s C-terminal tail and all subunits of the NDC80 complex (13). Their results employing optical tweezers demonstrated that the SKA complex itself on microtubule tips can bear load, and that in combination with the NDC80 complex, it strengthens the NDC80 complex-based tip attachments. Using size-exclusion chromatography, Huis et al. found that purified SKA and NDC80 complexes can interact independently of microtubule attachment. This direct interaction was also found to be dependent upon the C-terminal tail of SKA3. Purified SKA complexes containing truncated SKA3, lacking its C-terminal 104-412 amino acids corresponding to the entirety of the C-terminus starting just downstream of the intertwined junction of SKA subunits, failed to interact with the NDC80 complex (14). Additionally, they confirmed that phosphorylation of SKA3’s C-terminus by CDK1/Cyclin B promotes the SKA3-NDC80 complex interaction (14). These observations are consistent with Zhang et. al.’s findings that demonstrate phosphorylation of SKA3’s C-terminus by CDK1/Cyclin B, particularly at residues T358 and T360, are key for interaction of the SKA and NDC80 complexes (15).

Herein, we examine SKA3’s C-terminal tail utilizing tools predicting ordered and disordered regions or potential secondary protein structure discerned by homology amongst distantly related proteins. We identify a discreet region within the C-terminus that reveals more order and structure. We employ RNAi-mediated knockdown of endogenous SKA3 in combination with Doxycycline-facilitated induction of RNAi-resistant EGFP-SKA3 C-terminal variants and assess the variants’ abilities to support normal mitotic progression. We find that the SKA3 discreet region is critical for successful SKA complex-mediated mitotic progression. We generate a mutant with 3 sequential alanine substitutions within this critical region and find that it fails to support SKA complex function. We utilize AlphaFold3 to model interactions of SKA3’s C-terminus with the NDC80 complex and discover that the critical SKA3 C-terminal discreet region largely encompasses the residues within SKA3 that interact with the NDC80 complex. Since modeling predicted that the aforementioned SKA3 triple alanine substitution mutant would interfere with SKA3↔NDC80 complex interactions, we measured EGFP-SKA3 construct accumulation at kinetochores and found a reduction in kinetochore concentrated SKA when cells relied on this triple alanine substitution mutant. Using this modeling technique to inform directed mutagenesis we predicted that generation of a RNAi-resistant EGFP-SKA3 mutant with an alteration of just two amino acids within our proposed SKA3 NDC80-interaction motif, downstream of the location of the exchanged residues within our SKA3 triple alanine mutant, would also substantially disrupt the stability of the SKA3↔NDC80 complex interface and inhibit mitotic progression.

## Results

### Characterization of domains within SKA3’s C-terminus

Utilizing IUPRED3, a tool which predicts order within intrinsically unstructured proteins, we scanned the amino acid sequence of human SKA3 and found regions that were likely ordered (low scoring) or disordered (high scoring) (16). In agreement with studies characterizing the structure of the SKA complex, these predictions indicated that the first 100 amino acids of SKA3, essential to form the SKA complex by interacting with SKA1 and SKA2 subunits, are more ordered and thus likely to form a stable, fixed 3D structure. However, the C-terminal amino acids beyond amino acid 100 were predicted to be largely disordered with few extended regions receiving stably low scores indicating structure. We then used HHPred, employing homology detection and structure prediction by HMM (<u>H</u>idden <u>M</u>arkoff <u>M</u>odels)-HMM comparison, to further narrow our exploration of SKA3’s C-terminus. HHPred suggested secondary structure within the amino acid range 365 through 393, a section that also encompassed an IUPRED3-predicted region trending toward ordered structure (Figure 1A, Supplementary Figure 3, (17)).

**Figure 1.**
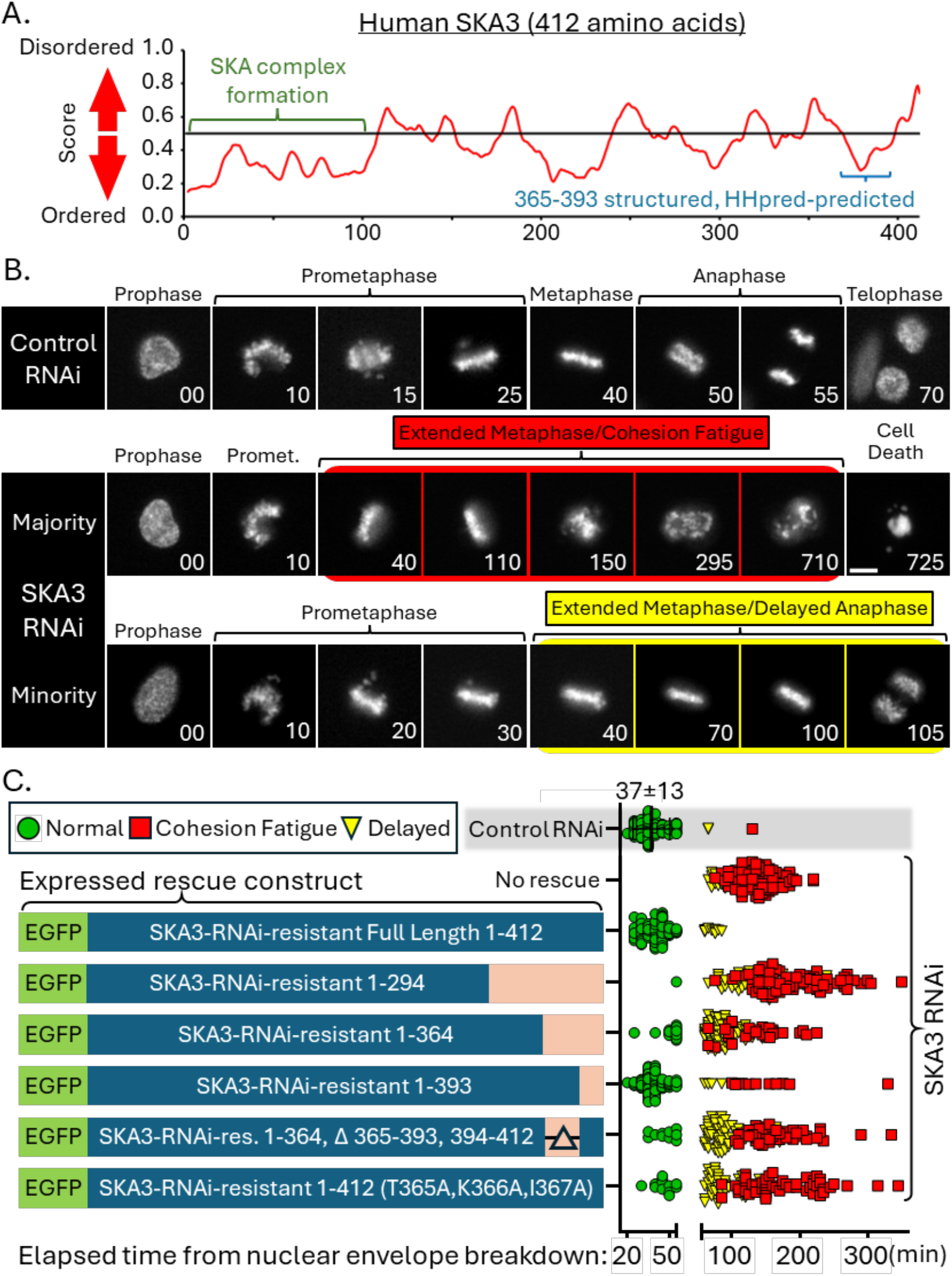
Human SKA3 ordered and disordered structural predictions and the mitotic fates of control and SKA3-depleted HeLa cells. (A) The 412-residue human SKA3 protein’s C-terminal 102-412 amino acid tail is predicted to be largely disordered by IUPRED3 and HHpred, albeit with a small section inclusive of amino acids 365-393 that both predictive tools suggest may be ordered and contain secondary structure. (B) Chromatin fluorescence image panels from HeLa live-cell time lapse microscopy comparing mitotic fates of control cells or those targeted for SKA3 depletion via RNAi. (C) Illustration of RNAi-rescue live cell assays categorizing fates of cells depleted of endogenous SKA3 via RNAi and rescued by induction of RNAi-resistant EGFP-SKA3 variants. Control RNAi data indicates the mean elapsed time duration from NEB to anaphase onset ± standard deviation.

To assess the ability mutant SKA3 constructs with varying lengths and C-terminal composition changes to rescue mitotic progression in the absence of endogenous SKA3, we created a series of stable isogenic HeLa cell lines expressing Doxycycline-inducible RNAi-resistant EGFP (<u>E</u>nhanced <u>G</u>reen <u>F</u>luorescent <u>P</u>rotein)-SKA3 constructs. The parental cell HeLa cell line lacking inducible constructs, termed HeLa Flp-In T-REx, was generated using the HeLa Flp-In T-REx system (Thermo Fisher Scientific, Catalog No. R71407).

Time-lapse live cell fluorescence microscopy tracking chromatin revealed that when transfected with control, non-targeting siRNA duplexes, parental HeLa Flp-In T-REx cells competently completed mitosis. Cells typically proceeded unabated through mitosis traversing from NEB (<u>N</u>uclear <u>E</u>nvelope <u>B</u>reakdown), the start of prometaphase, to anaphase at 37 minutes ± 13 minutes (Figure 1B, C). In contrast, when transfected with siRNA duplexes targeting endogenous SKA3, HeLa Flp-In T-REx cells failed to progress normally through mitosis and usually suffered one of two fates. The majority (75%) experienced an extended delay with chromosomes aligned at the metaphase plate followed by cohesion fatigue, which is asynchronous separation of chromatids induced by spindle pulling forces on chromosomes arrested at metaphase (18). The other 25% delayed their transition from metaphase to anaphase longer than the sum of the control’s average duration plus 1.5 times this average’s standard deviation and were categorized as “delayed” (Herein, delayed > 55 minutes, Figure 1B, C). Chromosome congression to the metaphase plate occurred normally in SKA depleted cells as previously reported (4, 5).

Doxycycline-induced expression of full-length RNAi-resistant EGFP-SKA3 largely rescued mitotic progression when endogenous SKA3 was targeted by RNAi, with 89% completing all mitotic stages in a timely manner while just 11% delayed at metaphase prior to initiating anaphase. But expression of RNAi-resistant truncated EGFP-SKA3 1-294, lacking amino acids 295-412 that correspond to roughly half of the unstructured C-terminus, poorly rescued mitotic progression with 73% arresting at metaphase and undergoing cohesion fatigue, 26% showing delayed anaphase onset, and 1% completing all mitotic stages in a timely manner. This truncated SKA3 lacks residues pT358 and pT360 known to be phosphorylated by CDK1/Cyclin B, crucial in promoting interaction with the NDC80 complex, as well as lacking the C-terminal region of interest spanning residues 365-393. However, expression of RNAi-resistant EGFP-SKA3 composed of amino acids 1-364, containing these important phospho-residues but lacking our region of interest also showed severe mitotic defects. In this condition, 86% of mitotic cells failed to progress normally through mitosis with 28% undergoing cohesion fatigue, 58% showing metaphase delay, and 14% exhibiting normal mitosis. The extreme C-terminal amino acids within human SKA3, amino acids 394 through 412, are highly conserved throughout the animal kingdom (Supplemental Figure 1). But expression of RNAi-resistant EGFP-SKA3 lacking these extreme C-terminal amino acids rescued mitotic progression nearly as well as the full-length construct, with 86% completing mitosis normally, 10% experiencing cohesion fatigue, and 4% delayed. Expression of an RNAi-resistant “Delta” EGFP-SKA3 construct lacking amino acids 365-393, our identified region of interest, but possessing the extreme c-terminal amino acids 394-412 fused after position 364, was as equally poor at rescuing the depletion of endogenous SKA3 as the truncated SKA3 construct lacking all amino acids post position 364. Expression of this construct in Ska3-depleted cells resulted in 44% of mitotic cells undergoing cohesion fatigue, 51% delaying at metaphase, and 5% progressing normally through mitosis. These data revealed that the SKA3 region encompassing amino acids 365-393 is critical for SKA complex-mediated mitotic progression.

To disrupt the structure within the 365-393 amino acid C-terminal SKA3 region critical for SKA complex function, we created an alanine substitution mutant by altering the first three amino acids within this putative functional motif. Expression of this RNAi-resistant EGFP-SKA3(T365A/K366A/I367A) mutant was unable to rescue the lack of endogenous SKA3 when assayed in our construct-inducible HeLa Flp-In T-REx system (Fig 1 B). The EGFP-SKA3(T365A/K366A/I367A) mutant construct’s inability to rescue normal mitotic progression was similar to that of the Delta mutant, with 53% exhibiting cohesion fatigue, 39% delayed, and 7% of cells proceeding normally through mitosis (Fig 1C). These data confirm that a key SKA3 functional element lies within SKA3’s C-terminal 365-393 region. When endogenous SKA3 was targeted by RNAi in our HeLa Flp-In T-REx cell lines, the Delta and T365A/K366A/I367A expressed RNAi-resistant EGFP-SKA3 constructs were capable of concentrating upon the mitotic spindle and kinetochores. This suggests that the inability of these truncated or mutant constructs to rescue mitotic progression was not likely due to gross mis-localization of the SKA complex (sup fig 2).

**Figure 2.**
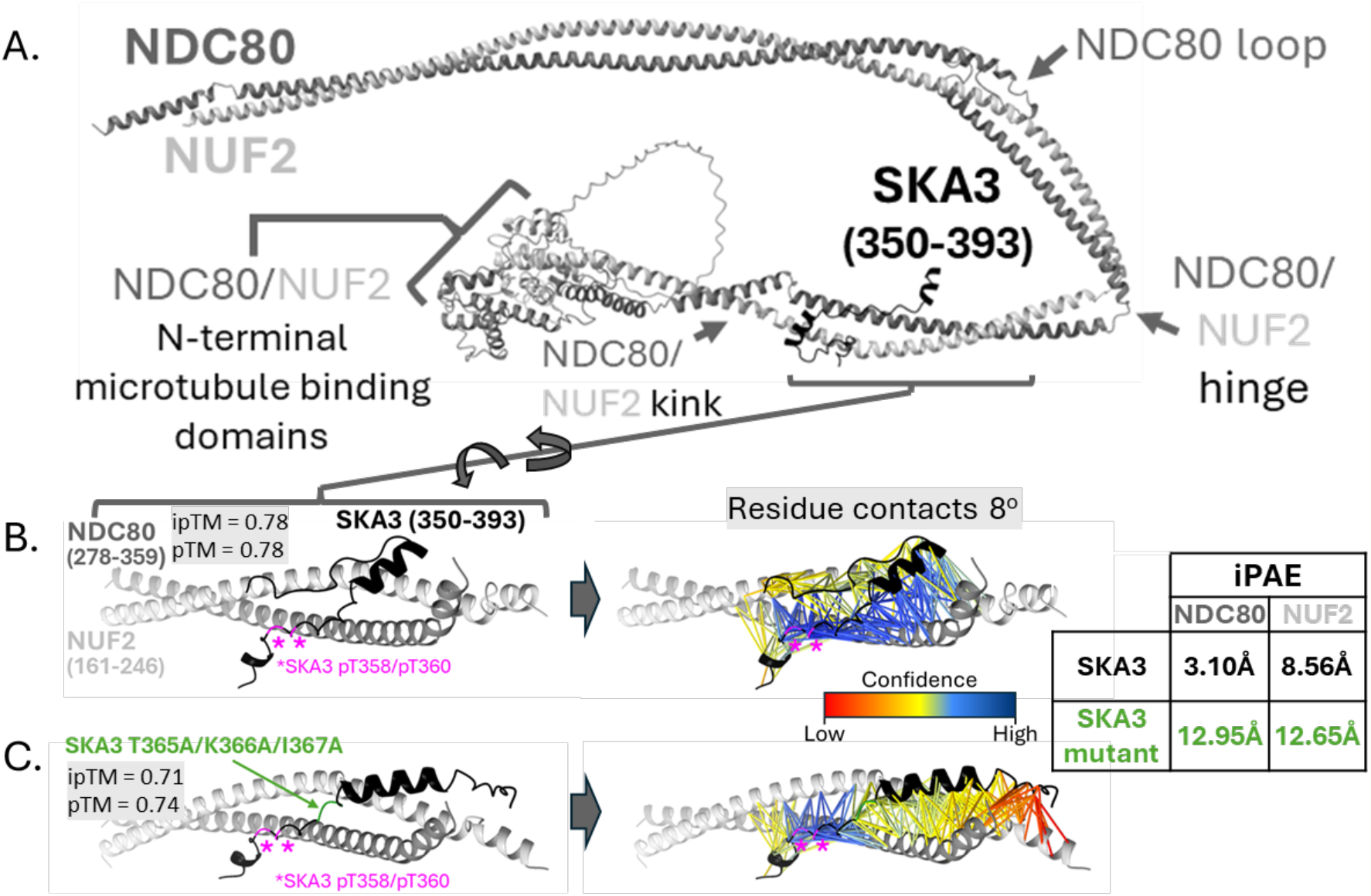
AlphaFold3-predicted structural protein models of NDC80 complex members NDC80 and NUF2 with SKA3’s C-terminal residues 350-393. (A) SKA3 C-terminal residues 350-393 interact with the coiled-coils of full length NDC80 and NUF2 proteins between a kink and the hinge, downstream of the microtubule domains. (B) Zoomed models of the SKA3 fragment interacting with fragments of the NDC80/NUF2 coiled coil. ipTM (interface Predicted Template Modelling score) and pTM (predicted template modelling score) confidence metrics are shown. Residue contacts of PAE ≤8Å with colorimetric confidence scale and iPAE (interface predicted alignment error) are reported. Post-translationally modified phosphorylated pT358 and pT360 SKA3 residues are magenta. Bar = 5 µm. (C) SKA3 T365A/K366A/I367A fragment modeled with NDC80/NUF2 fragment. Confidence metrics, colorimetric confidence scale, and iPAE are shown.

### Modeling the SKA3↔NDC80 complex interaction

Since interactions between SKA and NDC80 complexes mediated by SKA3 had been reported, we employed the open-source artificial intelligence system known as AlphaFold to predict where SKA3 might interact with the NDC80 complex and to determine if our SKA3 C-terminal mutant constructs would likely interfere with this interaction. AlphaFold version 3, developed by Google and DeepMind, predicts three-dimensional structures of proteins from their primary amino acid sequences and allows for modeling with inclusion of post translational modifications such as phosphorylated residues (19).

Directed queries predicted a model wherein a relatively short region of SKA3’s C-terminus interacts with both NDC80 and NUF2. This interaction is within a NDC80/NUF2 coiled-coil section downstream of the N-terminal microtubule binding globular heads but upstream of both a previously described NDC80/NUF2 “hinge” that provides apparent flexibility within the NDC80 complex (22). Additionally, our model predicts a “kink” in the NDC80/NUF2 coiled-coil just upstream of the area where SKA3 is predicted to interact that may provide additional flexibility within the NDC80 complex structure (Figure 2A). Modeling the SKA3 350-393 amino acid C-terminal span phosphorylated at residues T358 and T360 returned high confidence contacts less than or equal to 8Å ranging across amino acids 285-331 of NDC80 and 176-220 of NUF2 (Figure 2B). These contacts overlap the SKA3 365-393 amino acid region critical for mitotic progression and SKA complex function that we’d discovered in live cell studies. In contrast, modeling the phosphorylated SKA3 350-393 segment with T365A/K366A/T367A substitutions reduced most confidence measurements of contacts ≤8Å between the NDC80/NUF2 coiled-coil and the mutant SKA3 to low or moderate (Figure 2C). Though the confidence measurements for amino acids surrounding the pT358 and pT360 residues remained high, the model suggested that the 3 sequential alanine substitutions of SKA3’s 365-367 residues would significantly compromise normal interactions between SKA3 and NDC80/NUF2. Calculating the interface predicted error alignment, or iPAE (also known as the average interaction predicted alignment error), summarizes the interaction confidence between two peptides. The lower the score the greater the confidence of a stably interacting and well-defined interface between peptides. The iPAE scores of the SKA3 350-393 peptide with the NDC80 285-331 span or NUF2 176-220 span were 3.10Å and 8.56Å respectively, within a range supporting high confidence of peptide interaction interfaces. However, the iPAE scores of the SKA3 350-393 T365A/K366A/T367A alterations with NDC80 285-331 and NUF2 176-220 were 12.95Å and 12.65Å respectively, indicating the relative positions of the peptides to one another was far less certain and of low confidence (Figure 2 B, C)

### SKA3 T365A/K366A/I367A fails to fully concentrate at kinetochores

We performed quantitative immunofluorescence assays to measure the accumulation of the SKA complex at kinetochores, using EGFP fluorescence as a proxy for SKA complex localization in cells where endogenous SKA3 had been targeted by RNAi but rescued by expression of RNAi-resistant EGFP-SKA3 constructs. We compared the full length EGFP-SKA3 construct with our poorly rescuing yet least sequence-disruptive RNAi-resistant EGFP-SKA3(T365A/K366A/I367A) mutant. As noted previously, maximal kinetochore concentration of the SKA complex occurs at metaphase. Therefore, we examined mitotic cells at this stage for quantification of kinetochore-concentrated EGFP-SKA3. Since fluorescence intensities of the EGFP tagged SKA3 constructs were relatively low in the fixed cell preparations, we amplified fluorescence signals of the EGFP constructs by inclusion of anti-GFP antibody within our assay (Fig 3A). To account for differences between sizes of individual kinetochores, we divided the kinetochore EGFP fluorescence values by the fluorescence of the kinetochore-proxy centromere proteins identified by CREST serum, reporting these data as EGFP fluorescence per unit centromere. CREST fluorescence signals defined the regions used for quantification of both centromeric and EGFP-SKA3 fluorescence.

**Figure 3.**
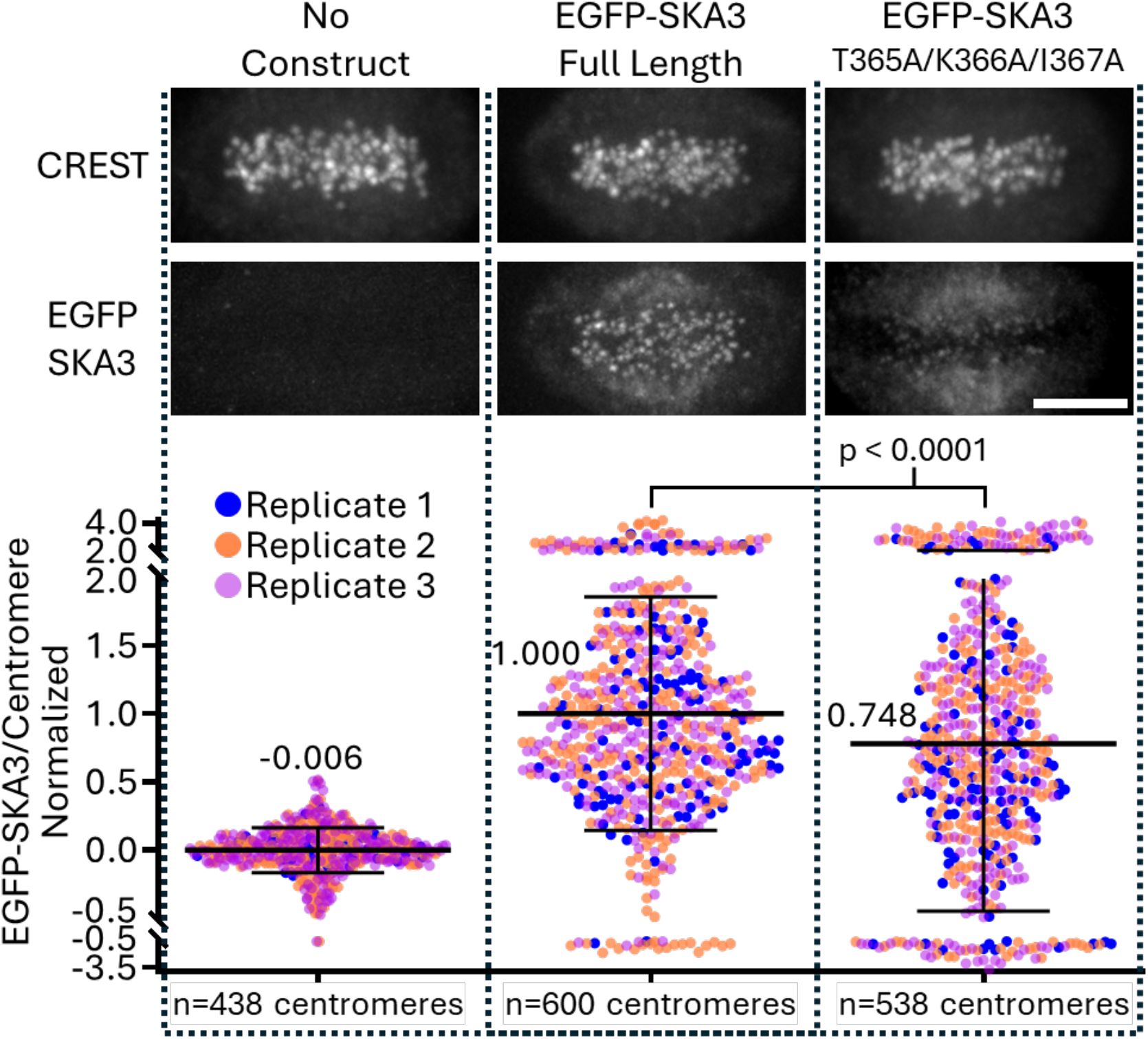
Quantitative immunofluorescence assay of metaphase cells showing normalized RNAi-resistant EGFP-SKA3 fluorescence per unit centromere in cells lacking endogenous SKA3 without rescue or rescued with expression of RNAi-resistant EGFP-SKA3 or EGFP-SKA3 T365A/K366A/I367A constructs. Results are compiled from 3 replicate experiments. The number of centromeres per category are indicated. Bar = 5 µm

The quantitative assay was replicated 3 times, with data compiled from all replicates. We set the positive control, metaphase cells expressing RNAi-resistant full length EGFP-SKA3 but lacking RNAi-targeted endogenous SKA3, to a normalized average value of 1.000 within each replicate (n=600 kinetochores from 42 cells, Fig 3B). The negative control, parental HeLa Flp-In T-REx cells lacking both RNAi-targeted endogenous SKA3 and an EGFP-inducible construct, confirmed that our assay results were consistent with an expected measurement for the absence of EGFP, returning an average value of 0.006 (n = 438 kinetochores from 37 cells, Fig 3B). In metaphase cells expressing the EGFP-SKA3(T365A/K366A/I367A) mutant construct but lacking endogenous SKA3, the average kinetochore associated EGFP-SKA3 construct measurement was 0.748 (n=538, kinetochores from 42 cells, Fig3B), a 25% decrease in the amount of accumulated SKA3 when compared to the non-mutated construct. Thus, this SKA3 (T365A/K366A/I367A) mutant fails to normally concentrate at kinetochores as predicted by the structural AlphaFold models indicating that the alanine substitutions of these three adjacent amino acids likely inhibit SKA3 interaction with the NDC80 complex.

### Model informed disruption of SKA3↔NDC80/NUF2 interaction by SKA3 C-terminal residue substitution

To further support the hypothesis that our SKA3 region of interest-truncated and sequential 3-alanine T365A/K366A/I367A mutant constructs do not support mitotic progression due to a disruption of the interaction between the SKA3 C-terminus and NDC80/NUF2 subunits, we created an additional SKA3 C-terminal construct with directed mutations informed by AlphaFold model predictions (Fig 4A). Substitution of two adjacent L (leucine) residues with D (Aspartate) residues within the SKA3 365-393 region at amino acid positions 374 and 375 predicted reduced binding of SKA3 to the NDC80 complex as compared to the endogenous SKA3 sequence. Most high confidence contacts between SKA3 and NDC80/NUF2 were reduced to moderate and low confidence when modeling the interaction with SKA3 L374D/L375D rather than the unaltered endogenous sequence. iPAE values between SKA3 and NDC80 or NUF2 peptides changed from high confidence values of 3.10Å and 8.56Å, respectively, to the low interface interaction confidence values of 13.97Å and 13.73Å when modeling the SKA3 L374D/L375D mutant (Figure 4A). After creating a Doxycycline-inducible RNAi-resistant EGFP-SKA3 (L374D/L375D) mutant HeLa Flp-In T-REx cell line, we assayed the ability of this EGFP-SKA3 construct to rescue mitotic progression. The majority of HeLa Flp-In T-REx cells transfected with non-targeting control siRNA duplexes completed mitosis in a timely manner. 96% of cells progressed normally through mitosis, 3% were delayed, and 1% delayed at metaphase long enough to induce cohesion fatigue. But cells transfected with siRNA duplexes targeting endogenous SKA3 and lacking expression of RNAi-resistant EGFP-SKA3 largely failed to progress through mitosis normally. In these cells 75% experienced cohesion fatigue, 22% underwent delayed anaphase, and 3% completed mitosis in a timely manner. Rescuing the lack of endogenous SKA3 with expression of non-mutant EGFP-SKA3 enabled 78% of cells to complete mitosis normally, with 18% delayed, and 5% experiencing cohesion fatigue. In contrast, expression of the EGFP-SKA3(L374D/L375D) mutant showed poor rescue with 27% completing mitosis normally, 42% delayed and 31% cohesion fatigue (Figure 4B).

**Figure 4.**
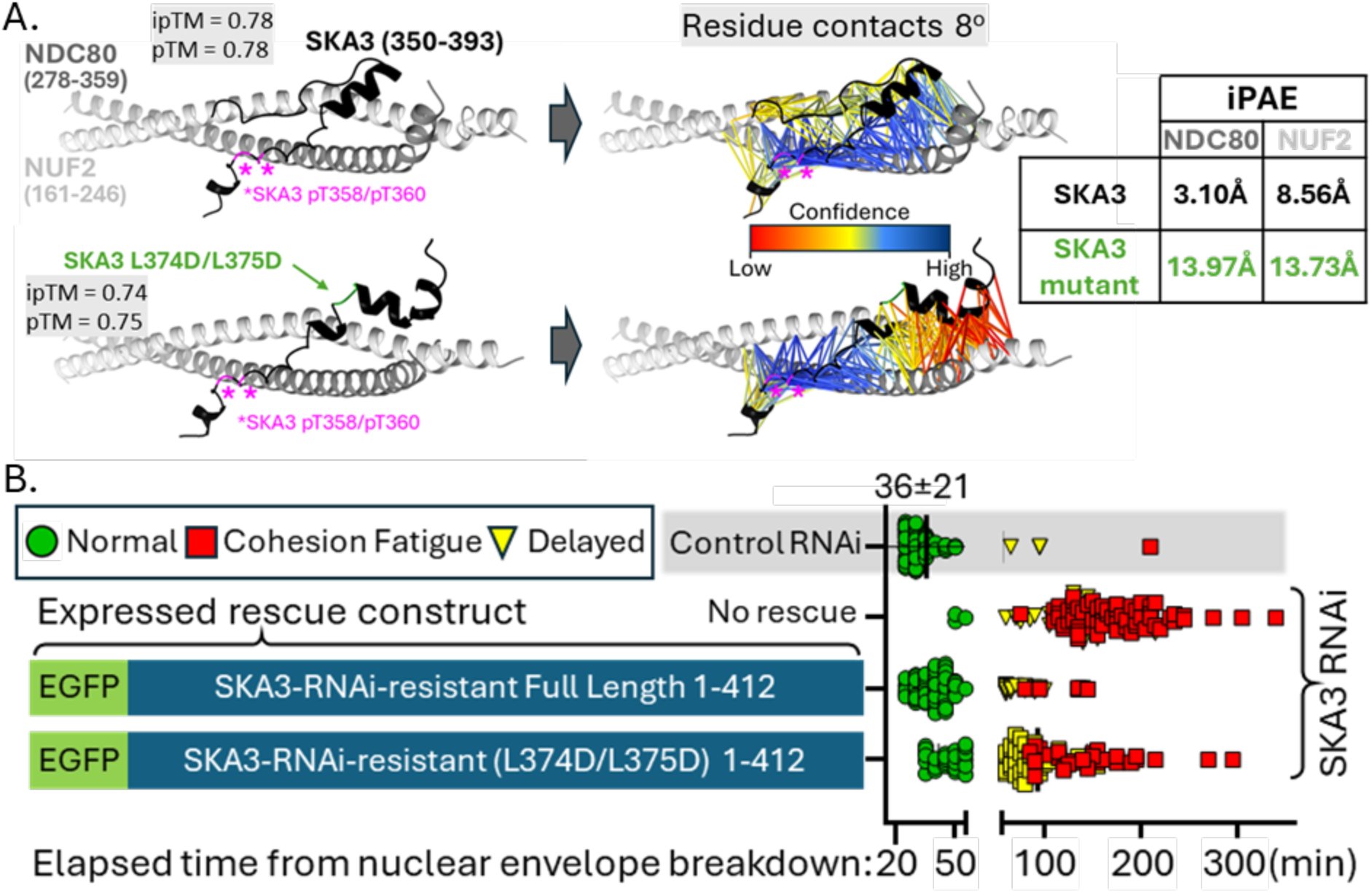
AlphaFold3-predicted structural protein models of NDC80/NUF2 coiled coil fragment with SKA3’s C-terminal residues 350-393 with or without model informed SKA3 L374D/L375D mutations. (A) ipTM and pTM confidence metrics are shown. Residue contacts of PAE ≤8Å with colorimetric confidence scale and iPAE are reported. Phosphorylated pT358 and pT360 SKA3 residues are magenta. (B) Illustration of RNAi-rescue live cell assays categorizing fates of cells depleted of endogenous SKA3 via RNAi and rescued by induction of RNAi-resistant EGFP-SKA3 variants. Control RNAi data indicates the mean elapsed time duration from NEB to anaphase onset ± standard deviation.

## DISCUSSION

How cooperation between SKA and NDC80 complexes ensure highly accurate chromosome segregation and successful mitotic progression has remained elusive. Herein, RNAi-rescue live cell time-lapse imaging assays provide insights into this cooperative interaction by defining a short region within the C-terminus of the third subunit of the SKA complex, SKA3, that is required for SKA complex function and successful mitotic progression. Guided in part by tools predicting order, disorder, or protein structure by homology detection, we created truncations of and mutations within the SKA3 C-terminus and assessed abilities of these variant SKA3 constructs to rescue loss of endogenous SKA3. Results of these live-cell mitotic progression assays indicated that a short region, spanning the 28 residues of human SKA3 from amino acids 365-393, was critical for normal mitotic progression. The region’s absence phenocopies a complete lack of SKA complex function.

Through structure prediction modeling we discovered that this SKA3 C-terminal 28 amino acid region lies within and constitutes the majority of an NDC80/NUF2 interaction module that includes additional upstream CDK1/Cyclin B phospho-threonine SKA3 residues at positions 358 and 360. Models generated without the post-translational phosphorylation of SKA3’s threonine 358 and 360 residues suggest a lower affinity interaction of SKA3 with the NDC80 complex (Supplementary Figure 4). This supports Zhang et al’s study demonstrating that phosphorylation of these SKA3 residues aids SKA complex function (15). Thus, our modeled and newly refined SKA3↔NDC80 complex interaction, largely inclusive of the important functional region of SKA3 that we’d defined by truncation and mutation, spans SKA3’s 41 C-terminal amino acids from residues 353-393. Per our model, SKA3’s C-terminal domain is predicted to interact with the NDC80 complex via the NDC80/NUF2 coiled-coil sections spanning NDC80 amino acids 285-331 and NUF2 amino acids 176-220.

Structure modeling was instructive as it informed the reasoning behind the inability of our SKA3 T365A/K366A/I367A mutant construct to rescue lack of endogenous SKA3, suggesting disruption of the SKA3↔NDC80 complex interaction. Predictively, fluorescence microscopy indicated that the SKA complex lacking endogenous SKA3 but containing our exogenous EGFP-SKA3 constructs could interact with microtubules and concentrate with microtubule ends at kinetochores. Although replacement of endogenous SKA3 with the EGFP-SKA3 C-terminal T365A/K366A/I367A mutant reduced the measured kinetochore concentration of SKA complex by just 25% when compared to the EGFP-SKA3 non-mutant construct, this altered SKA complex prevented mitotic progression and instigated cohesion fatigue, mirroring the SKA3-depletion phenotype. Since protein modeling predicted that this SKA3 T365A/K366A/I367A mutant disrupted the SKA-NDC80 complex interaction, we suggest that the reduction of SKA complex at kinetochores is reflective of the loss of SKA and NDC80 complex interactions. We surmise that the remaining intracellular concentrations of the mutant construct along spindle microtubules and at kinetochores are likely due to the microtubule and tip-tracking characteristics driven by SKA1’s C-terminal tubulin-interacting globular domain. While studies have shown that the loss of the entirety of SKA3’s C-terminus downstream of the N-terminal region intertwined with SKA1 and SKA3 causes a failure in SKA function and mitotic progression, this triple alanine mutant predicted to disrupt the modeled SKA3↔NDC80 complex interaction suggests that SKA relies upon this localized SKA3↔NDC80 interaction structural element for SKA-mediated mitotic progression via SKA’s role in NDC80 complex regulation.

Additionally, structure modeling enabled directed mutagenesis of SKA3 within the interaction motif that predicted disruption of the SKA3↔NDC80 complex interaction and by extension, inhibition of SKA complex function. This was realized within live cell mitotic progression assays demonstrating that our model inspired EGFP-SKA3 L374D/L375D mutant construct failed to rescue SKA function and normal mitotic progression. Collectively, these data indicate that the cooperative result of SKA and NDC80 complex interactions enabling spindle checkpoint silencing and anaphase onset after appropriate amphitelic attachments have been realized are likely dependent upon a roughly 40 amino-acid interaction motif near the extreme C-terminus of SKA3. Recently posted preprints (20, 21) of cryo-EM structural studies corroborate the location of the SKA3 and NDC80 complex interaction interface with the NDC80:NUF2 coiled-coil.

The location of SKA3’s interaction motif with the NDC80:NUF2 coiled coil supports hypotheses suggesting that the SKA complex may act to stabilize an extended structural conformation of the NDC80 complex reached when amphitelic spindle forces are achieved. This extended configuration is likened to a jackknife that has been extended and “locked” in an open position (22, 23). During prometaphase and prior to extension when microtubule connections are lateral or weak, NDC80 complexes are often bent, attributed largely to bending at the flexible region at the NDC80 hinge. This is termed the “closed” jackknife orientation. But as prometaphase transitions to metaphase and amphitelic microtubule-NDC80 complex interactions and tension increase, the NDC80 complex is extended as if a jackknife is opened and this configuration favors interaction with the SKA complex, likely via SKA3’s C-terminus. The jackknife hypothesis asserts that SKA complex then secures the NDC80 complex’s extended configuration in the locked or stabilized position. When the NDC80 complex is locked and favors persistent SKA complex interactions, SKA-mediated PP1 accumulation likely results in full PP1 kinetochore recruitment thereby opposing checkpoint kinase activities and promoting anaphase onset (8, 23, 24).

## Materials and Methods

### Generation of inducible EGFP-SKA3 fusion construct HeLa Flp-In T-REx cell lines

HeLa Flp-In T-REx cell lines (Thermo Fisher Scientific, Catalog No. R71407) were cultured in HyClone™ Dulbecco’s Modified Eagle’s Medium (DMEM), High glucose with L-glutamine and sodium pyruvate (Catalog # SH30243FS, Fisher Scientific, 4500 Turnberry Drive, Hanover Park, Illinois 60133-5491, United States) supplemented with 10% v/v Fetal Bovine Serum (Catalog# S1620-500, (USDA) ORIGIN, Biowest, 3135 Lakewood Ranch Blvd Suite 106 Bradenton, FL 34211-4957 United States), non-essential amino acids (HyClone™ Non-Essential Amino Acid (NEAA) Solution, 100X, Catalog# SH3023801, Fisher Scientific, 4500 Turnberry Drive, Hanover Park, Illinois 60133-5491, United States), 100 U/ml penicillin and 100 µg/ml streptomycin (HyClone™ Penicillin-Streptomycin Solution, 100X, 10000 U/mL Penicillin, 10000 µg/mL Streptomycin, Catalog # SV30010, Cytiva (Global Life Sciences Solutions USA LLC) 100 RESULTS WAY, MARLBOROUGH, Massachusetts 01752 United States), and 1 mM Hepes (Catalog# HOL06-6 × 100ML, Plant Cell Technology (formerly Caisson Labs) 836 South 100 East, Smithfield, UT 84335, USA).

HeLa Flp-In T-REx cell lines capable of expressing Enhanced Green Fluorescent Protein-Spindle and Kinetochore Associated subunit 3 fusion constructs were created by co-transfection of HeLa Flp-In T-REx cells with pOG44 and modified pcDNA.FRT.TO plasmids in a 9:1 molar ratio, respectively, with TransIT®-LT1 (Mirus Bio/Millipore Sigma, 5602 Research Park Blvd, Suite 210, Madison, WI 53719 USA). Transfected pOG44 expressed the Flp recombinase, required to “flip-in” the construct to the inducible genomic site from the modified EGFP-SKA3 fusion construct-containing pcDNA.FRT.TO plasmids. Hygromycin treatment at 200 µg/ml selected transfected cells that had appropriately incorporated the fusion construct.

### SKA3 RNAi interference and EGFP-SKA3 construct rescue

HeLa Flp-In T-REx cell lines were cultured overnight in wells of an 8-well glass-bottomed chambered coverslip (ibidi USA, Inc. 5525 Nobel Drive, Suite 130 Fitchburg, Wisconsin 5371, catalog# NC1754075) to a density covering 40% of the well’s surface area. Cells were transfected with control, non-targeting or SKA3-targeting siRNA duplexes at a final concentration of 25 nM using Lipofectamine RNAiMAX (Life Technologies) as per manufacturer recommendations. Control siRNA duplexes obtained from Dharmacon (a Revvity brand, 2650 Crescent Drive, Suite 100 Lafayette, Colorado 80026 USA catalog# D-001810-10). SKA3-targeting siRNA duplex sequence is 5’-GAUCGUACUUCGUUGGUUU-3’. For induction of EGFP-SKA3 constructs or mock treatment of control cells, Doxycycline was added to wells from an 11X stock in media to reach a final concentration of 1.5 ug/ml 6 hours post siRNA duplex transfection. Cells were cultured for an additional 22 to 24 hours prior to evaluation of mitotic progression (28-30 hours post siRNA duplex transfection).

### Timelapse live cell microscopy

Live timelapse fluorescence images were acquired using a Nikon TiE microscope equipped with a perfect focus system, a 20× (NA XX) air objective, DS-Qi2 camera (Nikon Instruments), and an Okolab environmental chamber providing a humidified environment with stable 5% CO_2_ and 37°C. DMEM-based media in cell culture wells was exchanged to DMEM-Fluorobrite (Gibco/Fisher Scientific, catalog # A1896701) supplemented with 10% Fetal Bovine Serum, penicillin, streptomycin, 2 mM L-Glutamine, 1 mM Sodium Pyruvate containing 500 nM sir-DNA and 10 uM Verapamil (Cytoskeleton, catalog # CY-SC007) or SPY650-DNA (Cytoskeleton, catalog # CY-SC501) with or without 1.5 µg/ml Doxycycline as indicated ∼ 60 minutes prior to image acquisition. This media exchange bathed the cells in a low fluorescence DMEM-based media and allowed cells to absorb the SiR-DNA or SPY650 cell-permeable and non-toxic fluorescent DNA dyes. Fluorophores were excited and detected using a CY5 filter set (Nikon, catalog# 96366, Excitation: 620/60nm (590-650nm), Emission: 700/75nm (663-738nm)). Acquisition intervals were every 5 minutes over an 18-to-24-hour period.

### Immunofluorescence assays

HeLa Flp-In T-REx cell lines were cultured on 22 × 22 mm number 1.5 sterilized glass coverslips. Cells were transfected with SKA3-targeting siRNA duplexes at a final concentration of 25 nM using Lipofectamine RNAiMAX (Life Technologies) as per manufacturer recommendations. 8 hours post transfection, Doxycycline was added at 2 µg/ml. 30 hours post transfection, cells were fixed by immersion for 15 minutes at room temperature in 1.5% Formaldehyde, PHEM (60 mM Pipes, pH 6.9, 25 mM Hepes, 10 mM EGTA, 4 mM MgSO_4_, pH 6.9), 0.5% Triton X-100, 1:1000 dilution Protease Inhibitor Cocktail (Sigma, catalog# P8340). After rinsing in MBST (Mops Buffered Saline (10 mM MOPS, 150 mM NaCl, pH 7.4) with 0.05% Tween 20), coverslips were incubated for 45 minutes in a solution containing 20% boiled normal goat serum (BNGS) and MBST to block non-specific protein binding sites. After blocking, coverslips were incubated for 2 hours in a 5% BNGS/MBST solution with primary antibodies. Primary antibodies were as follows: Rabbit anti-GFP antibody at 0.5 µg/ml (ChromoTek, catalog# PABG1); YL1/2 Rat anti-Tubulin alpha antibody at 1:1000 dilution (YL1/2, AbD Serotec); Nuclear ANA-Centromere Autoantibody (CREST) (Cortex Biochem, catalog# CS1058) at 1:1000 dilution. After washing 3 × 4 minutes in 2 ml MBST, coverslips were incubated for 2 hours in a 5% BNGS/MBST solution with secondary antibodies. Secondary antibodies were as follows: Alexa Fluor® 488 AffiniPure® Goat Anti-Rabbit IgG (H+L) (min X Hu, Ms, Rat Sr Prot) at 0.5 µg/ml (Jackson ImmunoResearch Laboratories, Inc., catalog# 111-545-144); Cy3-AffiniPure Goat Anti-Rat IgG (H+L) (min X Hu,Bov,Hrs,Rb Sr Prot) at 0.33 µg/ml (Jackson ImmunoResearch Laboratories, Inc., catalog#112-165-143); Alexa Fluor® 647 AffiniPure® Goat Anti-Human IgG (H+L) at 0.33 µg/ml (Jackson ImmunoResearch Laboratories, Inc., catalog# 109-605-003). Coverslips were washed 3 × 4 minutes in 2 ml MBST, then stained with DAPI (4′,6-diamidino-2-phenylindole) at 100 ng/ml in water for 3 minutes. After washing 3 minutes in water, followed by 3 rinses in water, the coverslips were mounted on slides using VECTASHIELD Vibrance® Antifade Mounting Medium (Vector Laboratories, catalog# H-1700).

### Confocal microscopy and quantitative fluorescence measurements

Confocal microscopy was employed to capture immunofluorescence assay images using a Slidebook (Intelligent Imaging Innovations)-controlled Zeiss AxioObserver with a 100 × 1.4X NA objective and a Yokogawa confocal scanning unit (CSU22) coupled to a Hamamatsu Orca-Flash 4.0 LT camera (C11440). Images acquired through Z with 0.28 µm steps. Quantitative measurements were obtained using Metamorph (Version 7.8.12.0, Molecular Devices, LLC) employing the method previously described by King et. al. (25). Punctate dots on paired sister chromatids from Nuclear ANA/CREST serum fluorescence were used as a proxy for kinetochore location. Concentric circular regions surrounding non-overlapping Nuclear ANA/CREST fluorescence puncta were selected from maximum Z-stack projections of metaphase cells. These regions were transferred to summed Z-stack projections of both Nuclear ANA/CREST and GFP fluorescence channels to calculate the Nuclear ANA/CREST and GFP fluorescence values within the kinetochore regions. GFP fluorescence per unit Nuclear ANA/CREST fluorescence for each kinetochore was calculated, then normalized to the mean of positive control.

### IUPRED3 and HHPRED predictive tools

IUPRED3 analysis of SKA3’s C-terminus was performed by submission of amino acid residues 1-412 within the tool using analysis type “IUPred3 long disorder” coupled with the advanced option “medium smoothing”. Graphical illustration of data was prepared using Microsoft Excel. HHPRED analysis of human SKA3’s C-terminus was performed by submission of amino acid residues 102-412 within the tool using default search parameters (Job-ID# 8509166).

### AlphaFold three-dimensional protein modeling

AlphaFold version 3 was utilized to model three-dimensional protein interactions between members of the human NDC80 and SKA complexes. Model illustrations were prepared using images derived from ChimeraX (Version 1.10.1). Shown in figure 2A is a model of full length NDC80 and NUF2 along with residues 350-393 of SKA3. Figures 2B-C and 4A contain models of NDC80 (residues 278-359), NUF2 (residues 161-246), and SKA3 (residues 350-393). ipTM (Interface Predicted Modeling) and pTM (Predicted Modeling) scores were calculated by AlphaFold. Models with truncated proteins permitted unfettered visualization of interactions between the truncated SKA3 and NDC80 or NUF2 molecules. In these models PAE (predicted error alignment) restricted to 8Å or less between SKA3 to NDC80 and SKA3 to NUF2 molecules are visualized using a colorimetric scale which correlates to the model-predicted confidence of accurate positioning of residues. PAE of 8Å or less between SKA3 and NDC80 and SKA3 and NUF2 molecules encompassed the entirety of the SKA3 350-393 molecule but were present only in residues ranging between 285-331 of NDC80 and 176-220 of NUF2. For calculation and comparison of predicted alignment error between normal SKA3 and mutated SKA3 models, average iPAE (Interface Predicted Alignment Error) was calculated by determining the average of PAE values from SKA3(350-393) to NDC80(285-331), the average of PAE values from NDC80(285-331) to SKA3(350-393), and then dividing the sum of these two averages by 2. Similarly, the average iPAE score for the interaction between SKA3 and NUF2 was calculated using SKA3(350-393) and NUF2(176-220) residues. Because phosphorylation frequently introduces significant local structural flexibility or multiple valid conformational states, AlphaFold3 returns a multitude of PAE values per residue within a single model for interactions involving the post-translationally modified phosphorylated SKA3 pT358 and pT360 residues. Prior to averaging PAE values within the matrix corresponding to the interacting residue spans within protein chains, the multitude of PAE corresponding to the individual phosphorylated residues were averaged and then the single averaged values per residue were used in the iPAE calculation. Residues without phosphorylation typically correspond to more highly conserved, single-stable conformations which leads AlphaFold3 to return a single, high-confidence (low PAE) result per residue. Thus, the iPAE values reported herein are determined by averages that equally weight each residue within the 3 interacting peptide chains.

## Acknowledgments

This manuscript is dedicated to the memory of Tim Pouland who played an important role in the research described. We thank the Center for Biological Data Sciences at the Oklahoma Medical Research Foundation for their assistance implementing AlphaFold. Research reported in this publication was supported by the National Institute of General Medical Sciences of the National Institutes of Health under award number 5R35GM126980. The content is solely the responsibility of the authors and does not necessarily represent the official views of the National Institutes of Health. Research reported here also benefitted from support by the McCasland Foundation.

---

**Supplementary Figure 1.**
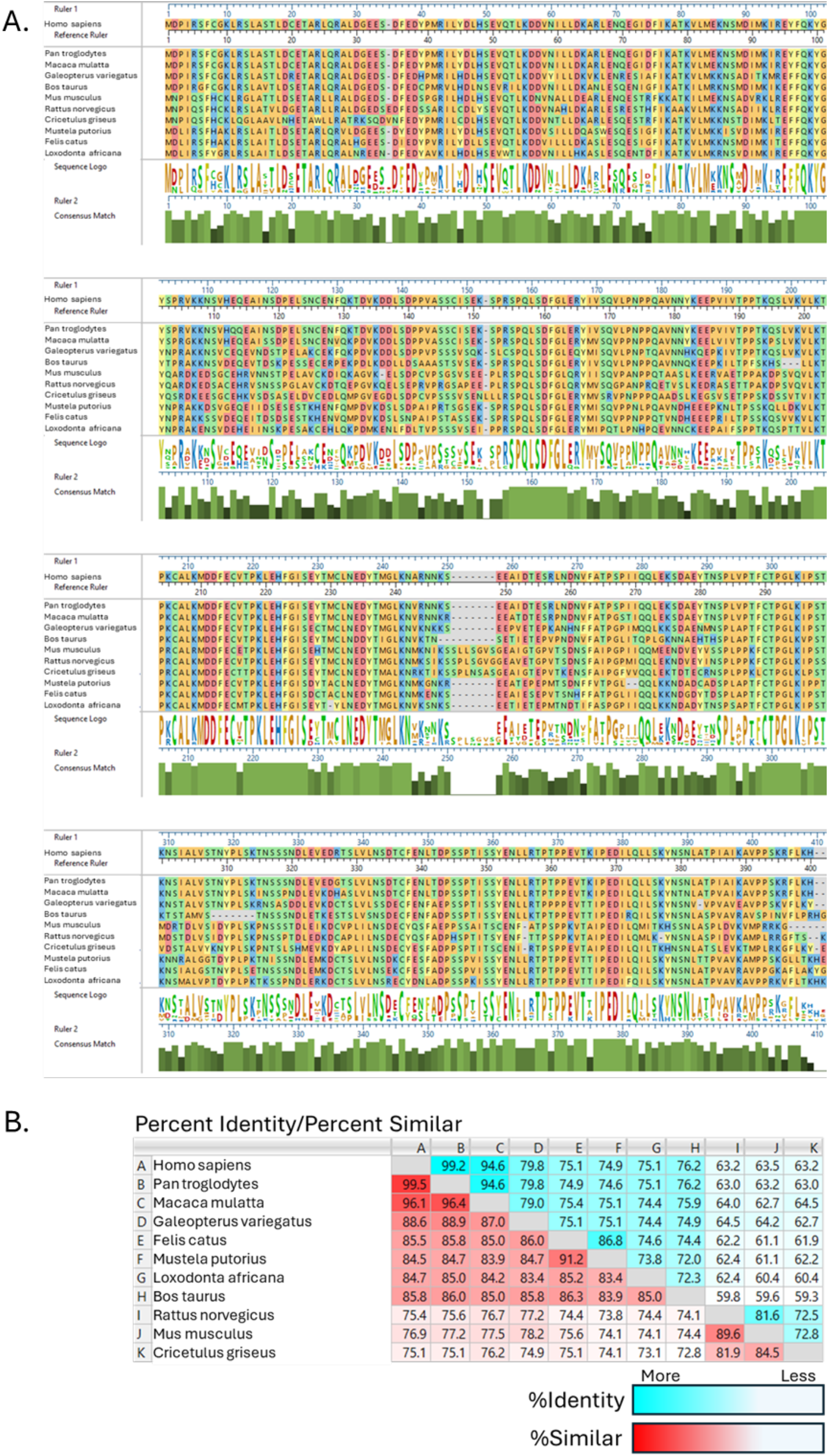
SKA3 protein alignments, identity, and similarity of various vertebrate species performed by DNASTAR MegAlignPro Version: 18.1.1 (8) using MUSCLE. (A) Residue depictions colored by chemistry. (B) Percent identity and similarity indicated numerically and colorimetrically by blue and red, respectively.

**Supplementary Figure 2.**
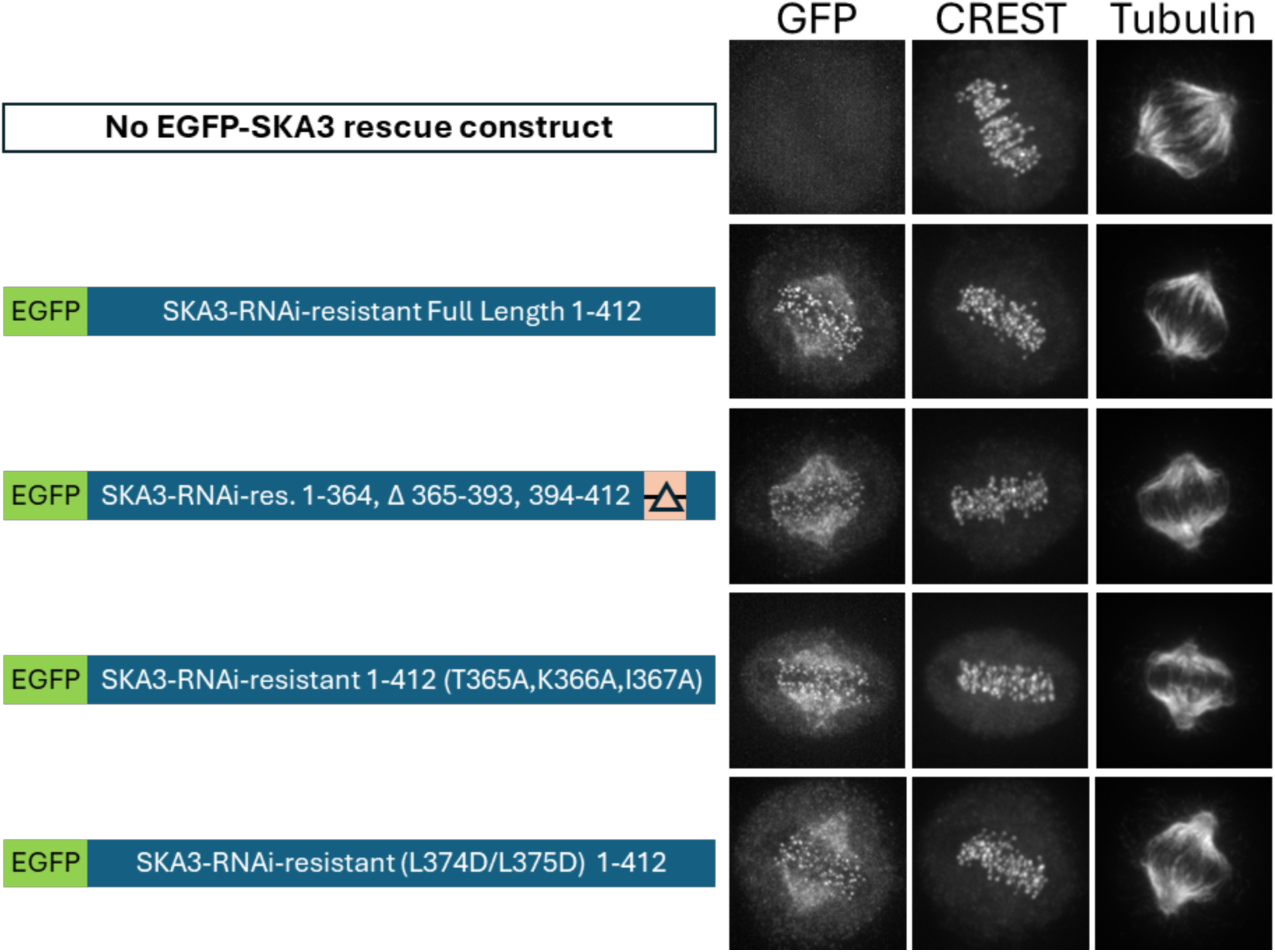
Immunofluorescence panels showing localization of expressed RNAi-resistant EGFP-SKA3 constructs via anti-GFP staining, CREST, and Tubulin in HeLa cells.

**Supplementary Figure 3.**
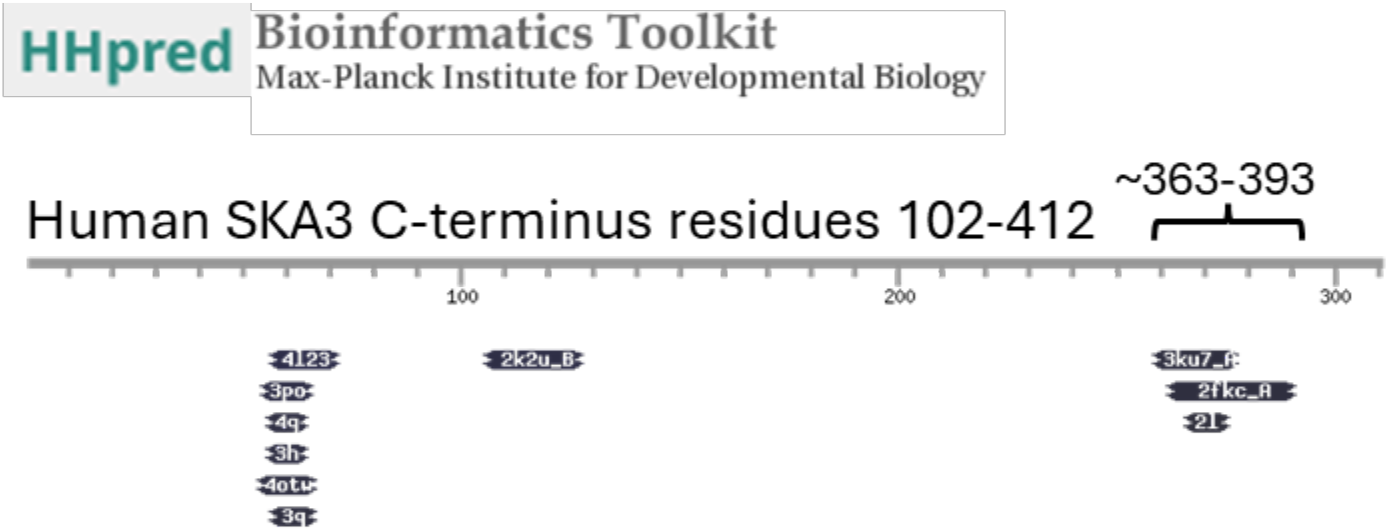
HHpred predictions showing potential structural similarity within the C-terminal residues 102-412 of human SKA3 protein with that of known but likely unrelated proteins suggesting ordered, secondary protein structure may exist within the largely disordered SKA3 C-terminus. A region from roughly amino acids 363 through 393 of SKA3 is accentuated.

**Supplementary Figure 4.**
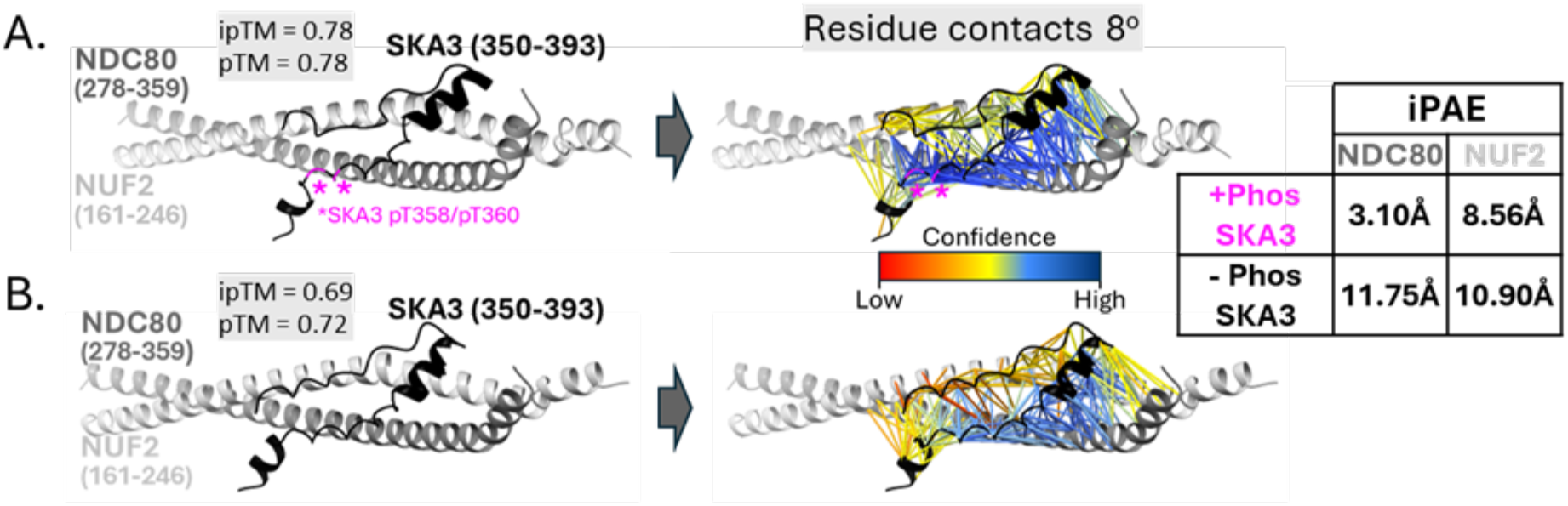
AlphaFold3-predicted structural protein models of NDC80/NUF2 coiled-coil fragment and SKA3’s C-terminal residues 350-393 with or without SKA3 phosphorylated residues. ipTM and pTM confidence metrics are shown. Residue contacts of PAE ≤8Å with colorimetric confidence scale and iPAE are reported. (A) Phosphorylated pT358 and pT360 SKA3 residues are magenta. (B) SKA3 modeled without phosphorylated residues.

